# Towards More Reliable Unsupervised Tissue Segmentation Via Integrating Mass Spectrometry Imaging and Hematoxylin-Erosin Stained Histopathological Image

**DOI:** 10.1101/2020.07.17.208025

**Authors:** Ang Guo, Zhiyu Chen, Fang Li, Wenbo Li, Qian Luo

## Abstract

Mass Spectrometry Imaging (MSI) provides a useful tool to divide a tissue section into sub-regions with similar molecular profiles, namely tissue segmentation. However, owing to the lack of ground truth, there is no reliable evaluation approach to assess the validity of unsupervised segmentation outcomes of MSI. We propose a novel solution grounded on a presumption that a segmentation is reliable if it can be reproduced using distinct bio-information extracted from independent sources. Specifically, besides molecular information from MSI data, we also obtain morphological information over a tissue section from its Hematoxylin-Erosin (H&E) stained histopathological image. MSI has high molecular specificity but low spatial resolving power, the H&E image has no molecular specificity but it can capture microscopic details of the tissue with a spatial resolution two magnitudes higher than MSI. The whole H&E image is split into an array of small patches, which correspond to the spatial pixels of MSI. A spectrum of informative morphological features is computed iteratively for each patch and spatial segmentation can be generated by clustering the patches based on their morphological similarities. Adjusted Mutual Information (AMI) score measures the degree of agreement between MSI-based and H&E image-based segmentation outcomes, which is defined by us as an objective and quantitative evaluation metric of segmentation validity. We investigated various candidate morphological features: a combination of Deep Convolution Neural Network (DCNN) features and handcrafted Threshold Adjacency Statistics (TAS) features finally stood out. The most appropriate number of tissue segments was also determined according to AMI score. Moreover, we introduced Co-Clustering algorithm to MSI data to simultaneously group m/z variables and spatial pixels, so potential biomarkers associated to each sub-region were discovered without the need of further analysis. Eventually, by integrating the segmentation outcomes based on MSI and H&E image data, the confidence level of the segment assignment was displayed for each pixel, which offered a much more informative and compelling way to present the segmentation results.

## 1. Introduction

Automatically segmenting a tissue sample into medically-relevant subregions is of particular interest in pathological and clinical applications[1][2][3]. To achieve this goal, certain type of biological imaging modality capable of characterizing the spatial heterogeneity of a tissue must be used. Mass Spectrometry Imaging (MSI) is a rapidly rising molecular mapping technology[4][5]. A mass spectrometer ionizes chemical compounds, separates them according to their mass-to-charge ratios (m/z), and eventually generates a mass spectra consisting of m/z variables and their respective abundance. By scanning an ionization probe over a 2-D tissue section, MSI can simultaneously measure the in situ spatial distribution of different sorts of chemicals in a single experiment without any prior tagging of the molecular targets. The development of soft ionization methods such as Matrix Asisted Laser Desorption/Ionization [6] and Desorption Spray Ionization (DESI)[7] significantly expands the range of analyzable molecular species including but not limited to proteins, peptides, lipids, metabolites, and drugs[8][9][10][11]. Their spatial variations can be correlated with underlying anatomical structures, metabolic pathways, pathological phenotypes, and so on, which may lead to the discovery of new bio-markers or molecular interactions. For instance, DESI-MSI was used in [12] to delineate the boundary of tumor tissues and further divide them into three tumor sub-types according to their distinct molecular profiles. Therefore, MSI, which characterizes the spatial heterogeneity of a biological sample based on the molecular profiles (mass spectra) collected from various locations, has the potential as an automatic segmentation tool for tissues.

MSI-based tissue segmentation relies on an effective exploitation of rich molecular information in MSI-generated big data: MSI data comprise 10^4^ mass spectra along with their corresponding spatial locations on sample and each spectrum records 10^2^ *−* 10^4^ m/z variables along with their relative abundance. Unsupervised clustering analysis is one of the most commonly used data-mining methods to handle the high dimensionality and complexity nature of MSI data[1]. For a set of objects, clustering analysis separates them into several groups in such a way that objects in the same group (namely a cluster) are similar to each other than to those in other groups (clusters). In contrast to conventional MSI data analysis approach, unsupervised clustering doesn’t require manually selected ions or regions. Therefore, hidden data structure as well as unforeseen trend/correlation in the image or spectral domains can be explored, which provides an approach to fully exploit the label-free strength of MSI technique. Pixels (i.e. ion sampling locations) can be clustered based on their mass spectral similarity (i.e. similarity in terms of molecular profiles), which leads to a spatial segmentation of a tissue section along image domain. Labeling each pixel by the color assigned to its cluster id provides an overview of the variation of molecular content of a MSI dataset. However, a variety of factors may affect the final segmentation result: noise or instabilities in the data acquisition process, the choices of clustering algorithm (such as hierarchical clustering, k-means, DBSCAN, etc.), key parameters for the clustering algorithm (such as a predefined number of clusters K in KMeans), and data preprocessing steps (such as dimension reduction methods[1]): PCA, pLSA, NMF, t-SNE, etc.). When no prior knowledge about the sample tissue (such as manual labeling by histology experts) is available, it is difficult to assess the quality of the segmentation and thus difficult to optimize the setup of the clustering pipeline in an objective and rigorous way. As a result, doubts may be raised concerning the robustness and reliability of segmentation outcomes using MSI.

We propose a novel solution to the above issue through an integration of multi-source bio-information. A tissue section is analyzed with regards to both its molecular and morphological variations, where molecular information is extracted with MSI and morphological information with HE staining histo-pathological image. The whole slide HE image is split into an array of small patches and each corresponds to a pixel of the MSI data. For each patch, a list of morphological features is computed to build a “morphological spectrum” using both handcrafted feature extractors (such as Haralick features to encode texture) and Deep Convolutional Neural Networks (DCNN, such as deep layers of VGG-16 to encode highly abstract vision concepts). Clustering analysis is performed on the mass spectra data as well as the morphological spectra data respectively and produces a pair of segmentation outcomes. Based on an intuitive assumption that a segmentation is more likely to be true if it can be well replicated by different bio-information modalities, Adjusted Mutual Information is used to gauge the resemblance of segmentation and the clustering pipeline can be improved by maximizing such resemblance. Moreover, we introduce co-clustering algorithm into MSI, which concurrently cluster both pixels and m/z variables. Salient bio-markers can therefore be determined for each sub-region without the hassle of additional statistical analysis. Finally, a confidence map is generated to visualize the reliability of the automatic segmentation by integrating the MSI-based and H&E-based outcomes.

## 2. Related works

### 2.1. Tissue segmentation with MSI

G.McCombie et al. pioneered the application of clustering analysis in MSI data to facilitate the definition of distinct spatial regions[13]. Two MALDI-MSI datasets acquired from a Alzheimer’s disease brain section and a complete rat head section respectively were analyzed by three different clustering algorithms (hierarchical clustering (HC), KMeans, and ISODATA) with Euclidean distance metrics. Prior to clustering, the authors used principal Component Analysis (PCA) to reduce the dimension and noise level of the datasets. Linear Discriminative Analysis (LDA) were employed to extract differential m/z variables between clusters. Deininger et al. analyzed gastric cancer tissue sections with HC and successfully divided tumorous and nontumorous tissues into separate clusters, which agreed reasonably well with histology [14]. To reduce nonbiological pixel-to-pixel variability and improve the smoothness of spatial segmentation maps, spatial or neighboring information in MSI data has been incorporated during clustering analysis. In [15], Alexandrov et al. applied an adaptive, edge-preserving denoising algorithm to each m/z channel before clustering. In a later work[16], the same group developed a spatially aware clustering approach, which defined a new distance between two pixels as a weighted sum of all spectral distances between counterpart neighbours surrounding each of them. The segmentation methods proposed in [15] and [16] were tested on two MSI datasets: a rat brain section and a section of a neuroendocrine tumor. More recently, Bemis et al[17] proposed a spatial shrunken centroids clustering method, which combines nearest shrunken centroid classification (a method for extracting subsets of informative features from highly multivariate data) with the above spatially aware clustering. Several biological tissues were segmented in [17], including a pig fetus cross-section and three rodent brain datasets.

Due to the lack of “ground truths”, none of the above works evaluated their tissue segmentation results in an objective and quantitative way. Important algorithm parameters, such as the number of clusters, were chosen either by expert[14][15], which was hard and subjective, or sophisticated internal criteria[**?**][18][**?**], which was purely based on information extracted from MSI data itself. A reliable evaluation method, where no human intervention is required but informative external reference is introduced, would therefore be ideal to assess the validity of unsupervised clustering results. Bio-information acquired for the same sample but with different bio-imaging modalities such as H&E image is supposedly correlated to that acquired by MSI and can be used as a trustworthy external reference.

### 2.2. Histopathology Image Analysis

Along with the digitization of histological tissue slides and the dramatic growth of computational power, a considerable amount of efforts has been devoted to the computer-aided analysis of pathology images[19]. A wide range of handcrafted features have proven their efficacy in quantitatively extracting histopathologically relevant information from a H&E image for automatic diagnosis/prognosis tasks. Interested readers are referred to [19] and [20] for more indepth reviews (features used in our work is briefly introduced in Section 3.3). More recently, Deep Learning (DL) based feature extractors have also been reported in the area of histopathology image analysis [**?**][21][22] (see Section 3.3.5 for more details). In [23], J.Baker et al. calculated a list of features for each tile of a whole slide image of brain tumor, divided the tumor tissue into sub-regions by unsupervisedly clustering the tiles, and selected an assembly of representative tiles of each sub-region for following supervised tumor type classification task. Our work follows a clustering pipeline similar to [23] for tissue segmentation but with a significantly distinct group of features.

### 2.3. The Integration of Multi-source Bio-information

There has been a growing interest in the integration of multimodal bioinformation data. In oncology, many subtypes of cancer can only be differentiated based on molecular differences and thus “molecular histology” has been developed which uses techniques such as immunohistochemistry (IHC) and fluorescence in situ hybridisation (FISH), as well as extraction-based molecular diagnostics such as gene sequencing together with classical histomorphology[24]. For instance, in [25] and [26], phenotypic traits (such as cell morphology) obtained by microscopic HE images were compared and combined with molecular traits (such as gene expression and mutations) obtained by high-throughput genomics in order to establish correspondences between them and build integrated feature embeddings for supervised learning tasks such as prognosis prediction.

In the field of MSI, it is also a well-established approach to use microscopic HE stain images together with MSI for combined histopathological and molecular analysis of tissue samples [27]. Conventionally, molecular profiles obtained by MSI are overlaid spatially with the HE images, which enables the correlation of proteins, peptides, metabolites, lipids, and drugs with histopathological features or tissue substructures. Staining has to be done after the MSI measurement of a tissue section to avoid mass spectral interference from stain chemicals. Moreover, a experimental and computational pipeline was proposed in [28] to overlay 3D bio-imaging modality magnetic resonance imaging (MRI) with 3D-MALDI imaging data of a mouse kidney, which opened new doors towards the integration of ex vivo and in vivo imaging data.

Recently, more sophisticated muti-modal data fusion methods have been developed to break the inherent lower-resolution limitation of MSI technique. In 2015, Raf et al [29] reported a data fusion framework for MSI and HE stain microscopy: a multivariate regression model was built to predict m/z variables of MSI using RGB variables (and their derivative variables such as saturation, hue, PCA, etc.) of a HE image, which improved the spatial resolution from 10 *µ*m to 330 nm. F.Vollnhals et al. [30] compared two pansharpening methods, Intensity–Hue–Saturation and Laplacian Pyramid, and demonstrated the latter was more robust for image fusion between Secondary Ion Mass Spectrometry and Electron Microscopy.

To date, there has been no report that compares tissue segmentation results based on MSI and HE image data, which offers a way to assess the validity of the segmentation outcome as well as a performance metric for evaluating different segmentation routines.

## 3. Methods

The core idea of our work is illustrated in Figure 1. Two biological imaging modalities, MSI and H&E staining microscopy, are used to respectively extract the spatially resolved chemical and morphological information of a tissue sample. So we obtain two hyperspectral data cubes: the first two dimensions (X-Y plane) of the data cubes correspond to 2D spatial locations, while the third dimension (Z-axis) of the first data cube corresponds to m/z variables of MSI data and that of the second one corresponds to morphological features computed from the H&E image. Using clustering analysis, the two hyperspectral data cubes generate two independent segmentation of the tissue and a comparison between them reveals whether a segmentation can be verified by distinct bio-information sources. Critical components of the whole segmentation pipeline are introduced in this section.

**Figure 1:**
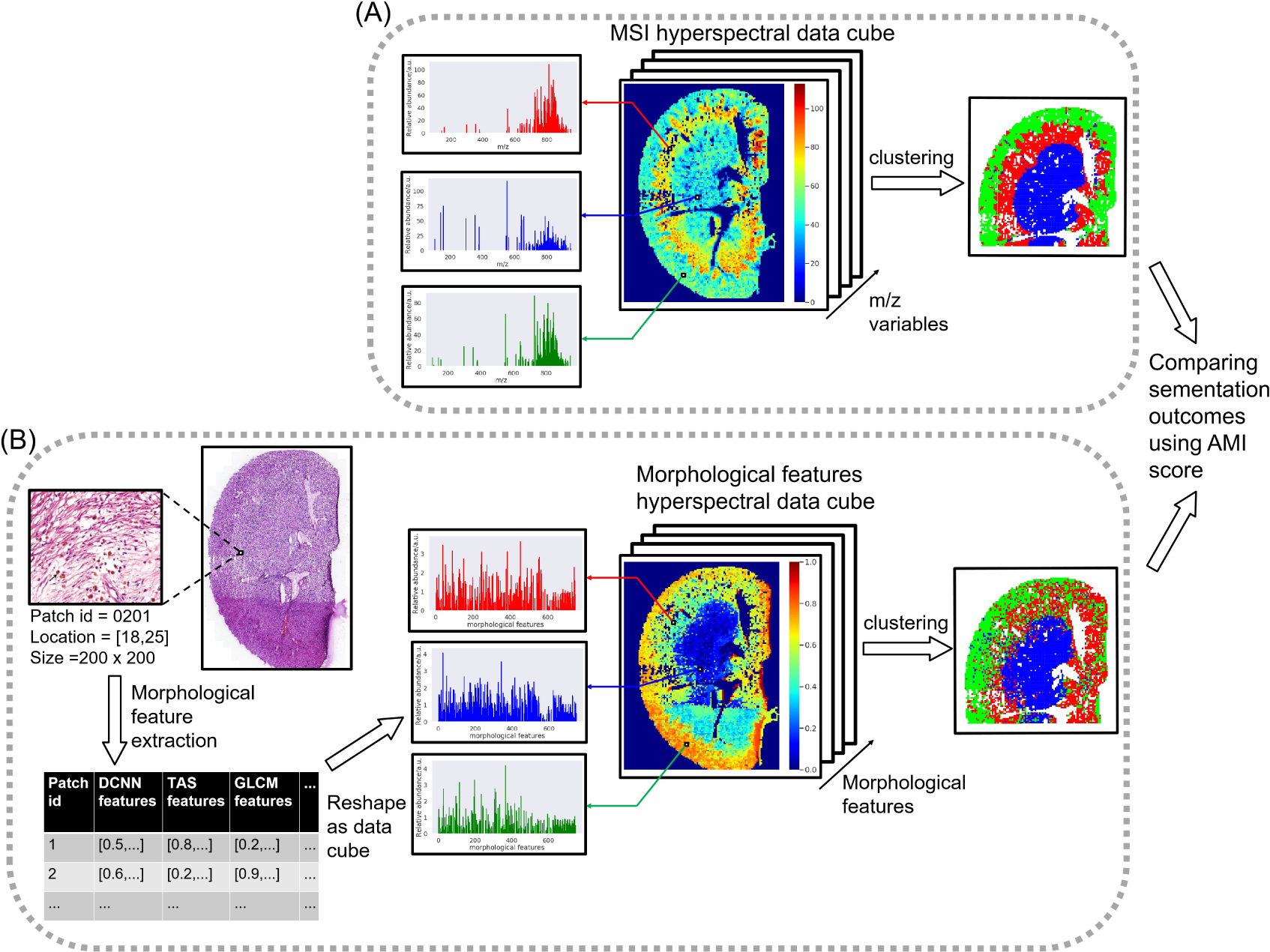
The overview of tissue segmentation based on (A) MSI data and (B) H&E staining microscopic data. In (A), mass spectra collected by MSI can be displayed as a 3D hyperspectral data cube. The X-Y plane holds the spatial information and the Z-axis represents the spectral information, namely the m/z variables. So each pixel of the X-Y plane corresponds to a mass spectrum (as shown by the red, blue, and green spectra on the left) and each channel of the Z-axis corresponds an intensity map of ions with certain m/z (as shown by the heat map in (A)). In (B), the high resolution H&E image is divided into an array of small patches and a set of quantitative morphological features are computed for each patch. So a similar 3D hyperspectral data cube can be generated as MSI data, with the only difference that the Z-axis corresponds to a spectrum of morphological features. Clustering on the basis of the similarity in the spectral domain result in a segmentation in the spatial domain. The segmentation outcomes based on two data cubes are correlated but not necessarily identical and comparison between them can reveal whether a segmentation can be verified by distinct bio-information sources.

### 3.1. Data Collection

We used tissue sections of a mouse kidney to illustrate the proposed segmentation method. The kidney sample was frozen-preserved at −80*°* before being cut into two adjacent slices by cryostatsectioning with thickness of 5 and 20 *µ* m, respectively. The thinner one went though standard Hematoxylin and Eosin (H&E) staining protocol, followed by digital pathology scanning with 20× magnification (i.e. 0.5 × 0.5 *µ* m pixel resolution). Raw MSI data was collected over the thicker one with a 100 × 100 *µ* m step size (or pixel resolution) using a DESI ion souce coupled to a Synapt G2-Si mass spectrometer (Waters, UK). A mass range of 200 - 1,000 m/z was covered with 60,150 mass bins over 13,608 pixels. It is worth noting that we assumed most structural features of the two adjacent slices were well conserved slice to slice and they could be thought identical.

### 3.2. Data Preprocessing

#### 3.2.1. H&E Staining Digital Pathology Image

By tissue detection, a binary mask was generated that delineated the area occupied by the kidney tissue in the digital pathology image. With the tissue mask as an input, Reinhard stain normalization method [31] was then used to transfer the colour characteristics of the tissue area to a desired standard with the intent to correct any staining or imaging variations. Both the preprocessing procedures described above were realized in python using HistomicsTK package [32].

#### 3.2.2. Mass Spectrometry Imaging

Raw MSI data was preprocessed through a sequence of standard operations: (1) Total-ion-count (TIC) normalization is widely used in literature [33][34] to project spectra of varying intensity onto a common intensity scale and therefore handle the experimentally introduced pixel-to-pixel variation of MSI data. The spectral profile of each pixel are scaled based on an assumption that the TIC collected on every pixel should be identical. (2) Spectral smoothing was obtained with a Gaussian kernel (window = 5 and standard deviation = window/4); (3) Baseline reduction algorithm interpolated a baseline from local minimas and subtracted it from the original spectral profile. (4) Peak picking (aka peak detection) identified meaningful m/z peaks by seeking local maxima above certain predefined signal-to-noise (SNR) threshold (SNR=6) in a sliding window (window width=5). In our case the adaptive noise was estimated by local mean absolute deviations (MAD).(5) Peak alignment, in order to eliminate tiny m/z-value shift due to instabilities of MS instruments, peaks with proximate m/z-values were matched given a tolerance threshold (200 ppm). (6) Peak binning, the intensity of a selected peak was represented with the sum of intensities between the two nearest local minima in both directions around its m/z-value. (7) Peak filtering determined the proportion of pixels where a peak was detected at a given m/z-value and only retained peaks with frequencies greater than 1%. Cardinal 2 package [35] in R was used to implement all the abovementioned procedures, which eventually output a MSI dataset consisting of a list of m/z-values for each picked peak (532 unique m/z-values), a list of the corresponding positions for each pixel (13,608 pixels), and a matrix of peak intensities (13, 608 × 532).

For MSI dataset, a binary mask was also produced to indicate pixels corresponding to the tissue sample (rather than background). Based on an assumption that there should be more large biological molecules residing in the tissue area than in the background, the sum of peak intensities between 500 and 1000 m/z were calculated for each pixel, and one would be tagged as ‘1’ (‘tissue’) if its sum was above a manually-set threshold and vice versa. To further reduce the set of m/z-values, t-tests were used to select ions that were significantly more abundant in the tissue than in the background. Consequently, 171 more informative peaks were retained for subsequent analysis.

### 3.3. Feature construction for H&E image

“Features” are a series of measurable properties to characterize an object in a data set. For MSI data set, it is rather straightforward that the m/z values are the features sufficient for extracting chemical composition information for each spatial pixel. However, constructing features to encode morphological information from H&E image isn’t as trivial. Traditional textural descriptors used to be most commonly found in the literature of digital pathology [20], but recently using deep convolutional neural networks (DCNN) as feature extractors is becoming increasingly popular in the era of Deep Learning. In our work, both conventional texture-related features and DCNN-extracted features were computed and combined for each spatial pixel. In this section, we briefly describe the essence of all the morphological features fed into the following clustering algorithm, including gray-level cooccurrence matrix (GLCM) [36], threshold adjacency statistics (TAS) [37], intensity statistics, as well as transfer learning based DCNN features [38].

#### 3.3.1. Gray-Level Co-Occurrence Matrices

GLCM are used to describe image texture[36]. Each element of GLCM is the probability that pixels of certain gray levels are spatially adjacent to each other. In our work, eight gray levels and four adjacency directions (horizontal, vertical, left and right diagonals) were used to construct the GLCM. 13 Haralick’s texture features were computed from GLCM [36]: angular second moment, contrast, correlation, sum of squares, variance, inverse difference moment, sum average, sum variance, sum entropy, entropy, difference variance, difference entropy, information measures of correlation 1, and information measures of correlation 2. Averaging in the four directions resulted in a final 13-D feature vector for each gray scale image.

#### 3.3.2. Threshold Adjacency Statistics

In TAS[37], 3 different threshold ranges are applied to the input image to create three binary images: [*µ* + *σ, µ* - *σ*], [*µ* - *σ*, 255], and [*µ*, 255], where *µ* is a threshold determined by the Otsu algorithm, and *σ* is the standard deviation of the above threshold pixels. A normalized histogram of 9 bins is generated for each binary image by counting the number of pixels surrounded by a given number (zero to 8) of white neighbours. Eventually TAS concatenates the above histogram with its bitwise negated version and outputs a 18-D feature vector.

#### 3.3.3. Intensity statistics

A list of intensity-derived features can be computed from an image, such as minimum, maximum, mean, median intensity of object pixels; difference between mean and median intensities; standard deviation, inter-quartile range, median absolute deviation, skewness and kurtosis of the intensities of pixels; energy and entropy of the intensity histogram of pixels. So, a total of 12 features was generated according to intensity statistics of each gray scale image.

#### 3.3.4. Nuclei Densities

Nuclei were detected and segmented by combining adaptive multi-scale LoG filter and local maximum clustering method. Then, 24 nuclei density related features were calculated including neighbor count within different radius and minimum distance to enclose count neighbors.

#### 3.3.5. Deep convolutional neural network

Last 8 years has seen a phenomenal success of DCNN in a variety of different image understanding tasks including the recognition, object detection, and semantic segmentation of images[39]. Unlike traditional approaches where images are characterized by domain-specific “handcrafted” features, DCNN takes a domain agnostic approach that combines both feature discovery and implementation to encode most useful properties of an image for the following tasks. Moreover, DCNN trained on large-scale computer vision data sets, such as ImageNet and Coco, have proven to be excellent, off-the-shelf feature extractors, capable of generating a set of generic features even for a distinct but reasonably relevant task. This is a widely accepted idea called “transfer learning”[38]. In our work, we used the VGG-16 network [40] pretrained using the well-known ImageNet database which contains 1.28 million images in 1,000 classes. After being resampled to match the 256 × 256 × 3 input size required by VGG-16, H&E images were forward propagated to the last convolutional layer of block 4. And the activations at that layer were 2-D globally average-pooled and extracted as 512-D feature vectors. The hierarchical structure of DCNN fully exploits the compositional nature of images, where higher-level features are obtained by combining lower-level ones: local edges assemble into motifs, motifs form parts, and parts form objects [39] [41]. Therefore, the features extracted by the deeper convolutional layer correspond to complex and abstract vision concepts, which are complementary to those handcrafted textural features. However, it is not self-evident to decide which layer is the most appropriate one for H&E image analysis. We compared clustering results using DCNN features extracted from different layers with that using MSI features and chose the one that resulted in the most similar segmentation.

To calculate the GLCM, TAS, and intensity statistics features for nuclei and cytoplasm separately, we deconvolved a RGB H&E image patch into two gray scale images that corresponded to the hematoxylin and erosin stains respectively. So the total number of morphological features was 2 × 13 + 2 × 18 + 2 × 12 + 512 = 598. The calculation of the GLCM and TAS features was implemented by the Mahotas package, intensity statistics by the HistomicsTK package, and DCNN by Keras.

### 3.4. Mapping between MSI pixels and H&E image patches

The pixel size of MSI was 200 times larger than that of H&E image, so each MSI pixel corresponded to a 200 × 200 patch of the whole slide image(*M*_*whole*_). We established an one-to-one mapping between a MSI pixel (*D*_*msi*_[*i, j*, :]) and a H&E image patch through the spatial registration technique: (1) the binary tissue mask of MSI data *M*_0_ was up-sampled by 200 times 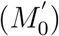 to have the same resolution as H&E image; (2) using 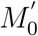 as fixed image and the binary tissue mask of H&E image *M*_1_ as moving image, an optimal Euler 2D transform matrix (*M*_*trans*_) was obtained with the gradient descent algorithm, which minimized the mean square metric between 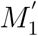 (*M*_1_ after transformation) and 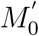; (3) applied *M*_*trans*_ to *M*_*whole*_ and output 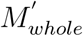.So 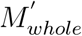 was in the same spatial coordinate system with 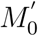, or in other words, the tissue area of 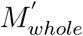 overlapped with that of 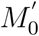; (4) consequently *D*_*msi*_[*i, j*, :] was mapped to H&E image patch 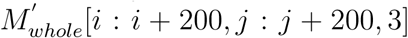. While iterating over every pixel, if it was labeled as tissue by *M*_0_ and no less than 90% of its corresponding H&E image patch was labeled tissue by 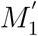, morphological features were calculated according to the image patch and stored along with its mass spectral features to form a new data frame: 9,999 rows corresponding to eligible pixles and 769 columns corresponding to 171 m/z values plus 598 morphological features. Spatial registration was realised by the SimpleITK library of python.

### 3.5. Clustering algorithms

Mining the sub-regions of a tissue section from its MSI data is achieved by clustering the pixels according to the similarity of their mass spectral profiles. Spectral co-clustering algorithm simultaneously groups the rows and columns of a data array, which is originally proposed to solve the documents-words co-clustering problem[42] but is also well suited to MSI data. The co-clustering task is modeled as a bipartite graph partitioning problem and solved by finding minimum normalized cut vertex partitions in the bipartite graph between rows and columns. Although finding a globally optimal partition is computationally prohibitive, it has been proved that the real relaxation of this NP-complete problem can be solved with the second left and right singular vectors of a dataset-derived matrix. More details can be found in [42]. One strength of spectral co-clustering over other clustering algorithms is its intuitive visualization and interpretation: by rearranging the MSI data matrix to make co-clusters contiguous, we can easily see that pixels are grouped together because certain m/z are significantly more abundant in their spectra and m/z variables are grouped because they have more abundance in the same bunch of pixels. Before co-clustering, the abundance was scaled for each m/z feature to have the same median and quantile range. Both the spectral co-clustering and scaling were implemented by scikit-learn library [43] in python. H&E image patches were clustered using the standard K-means algorithm in scikit-learn with cosine distance as multi-variable similarity metric.

## 4. Results

### 4.1. Optimizing segmentation pipelines

Based on an assumption that the segmentation of a tissue is more reliable if it can be corroborated by bio-information data from independent sources, we used Adjusted Mutual Information (AMI) score as evaluation metric to select features for morphological-features-based segmentation and to determine clustering algorithm for MSI-based segmentation. AMI score is a gauge to compare two sets of clustering results: perfect agreement gives an AMI score of 1.0 and random labeling gives an AMI score close to 0.0. Rooted in probability theory and information theory, AMI corrects the effect of agreement solely due to chance.

#### 4.1.1. Selecting DCNN output layer

As aforementioned, different convolution layers describe different levels of vision concepts: shallower layers tend to describe simpler concepts and deeper layers more complex ones. Since we use DCNN as morphological feature extractors, a natural question is which layer better fits our segmentation task. The pooling layers of the 3rd, 4th, and 5th blocks of the pretrained VGG16 were chosen as candidates and used to compute output arrays for each H&E image patch (block 5 is the last convolution block before the fully connected layers). After applying global average pooling on each output array, we constructed three feature vectors with a length of 256, 512, and 512 for the 3rd, 4th, and 5th blocks respectively. Then each feature was scaled to have zero median and unit variance across patches. The dimension of the feature vectors was reduced to 3 using UMAP algorithm before conventional KMeans clustering analysis. For MSI data, the abundance of each m/z variable was centered to its median along pixels and scaled according to an interquartile range 25% to 75%. Again, KMeans algorithm was used to cluster the MSI pixels. The AMI score was calculated and plotted against the number of clusters K which rose from 2 to 9. It is clear that block 4 prevails in all K values. A possible explanation is that those features extracted by block 5 may be too specific to the object recognition task of ImageNet and thus not quite relevant to our histopathological analysis, while block 3 features are perhaps too basic and thus uninformative. This conclusion is also consistent with [44], where models trained using deep inner layer features showed better accuracy in digital pathology classification tasks than those using last layers or shallow inner layers. We also tested another DCNN structure called Xception and its performance was considerably inferior to VGG16. Besides, there is a trend in Figure 2 that AMI score decreases in an asymptotic way as K increases, which indicates that the reliability of the unsupervised segmenation decreases as the number of clusters K increases, so K=2 or K=3 are probably the most appropriate number of sub-regions on the kidney tissue sample. This is in accordance with the anatomical structure of a kidney, which can be roughly segmented as pelvis, renal cortex, and renal medulla. The dependence of AMI scores on K are called “AMI score curves” thereafter.

**Figure 2:**
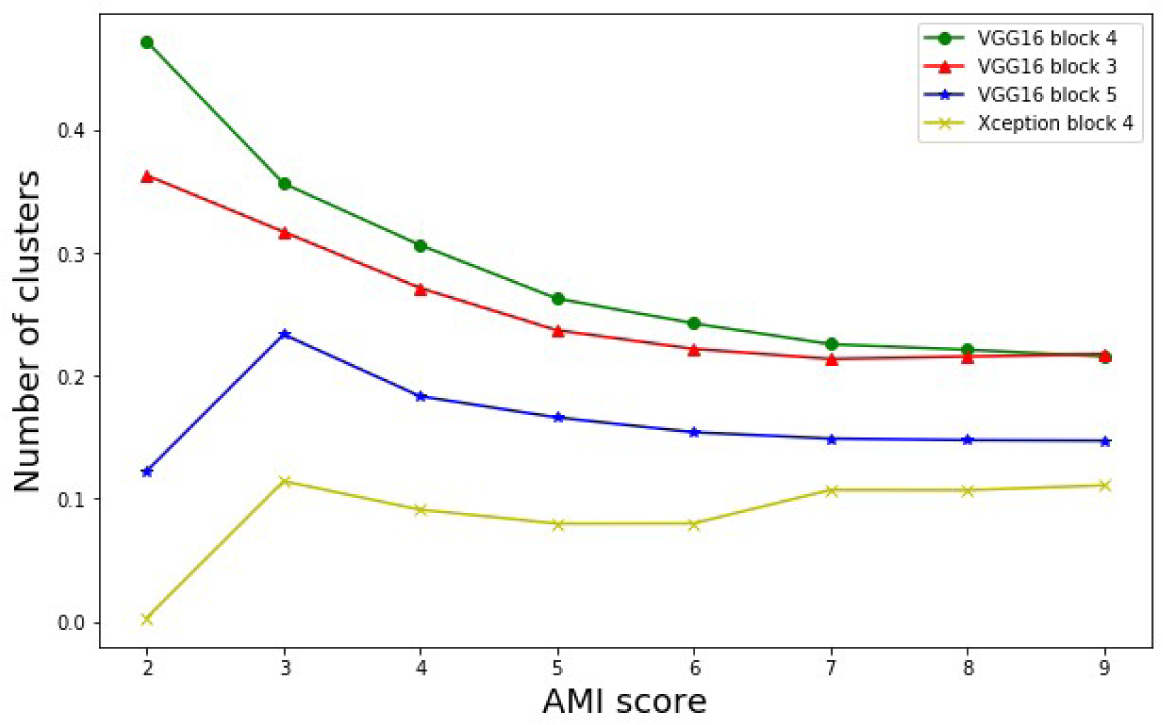
The dependence of AMI score on the number of clusters for 4 DCNN feature extractors. The pooling layer of the 4th block of VGG16 has shown the highest AMI score which means its segmentation outcome is in better agreement with the MSI-based one.

#### 4.1.2. Selecting handcrafted morphological features

Conventional handcrafted features have been widely used in the analysis of digital H&E image [23][20]. So, in order to dig as much useful morphological information as possible from a H&E image patch, a variety of handcrafted features was concatenated with DCNN features as the final morphological feature vector. Four sets of morphological features were investigated including Gray-Level Co-Occurrence Matrices (GLCM), Threshold Adjacency Statistics (TAS), Intensity statistics (IS), and Nuclei Densities (ND) (see section 3.3 for more details). Clustering analysis followed the same procedures as 4.1.1. As shown in Figure 3, AIM scores are mostly comparable for different combinations of feature sets. Since the combination of DCNN and TAS outperforms DCNN-only at K=2 and K=3 (which correspond to the most reliable segmentation outcomes), we choose DCNN and TAS to build the final morphological feature set. TAS uses pixel-wise statistics to describe image texture, which is complementary to the relatively complex vision concepts described by the deep layers of DCNN.

**Figure 3:**
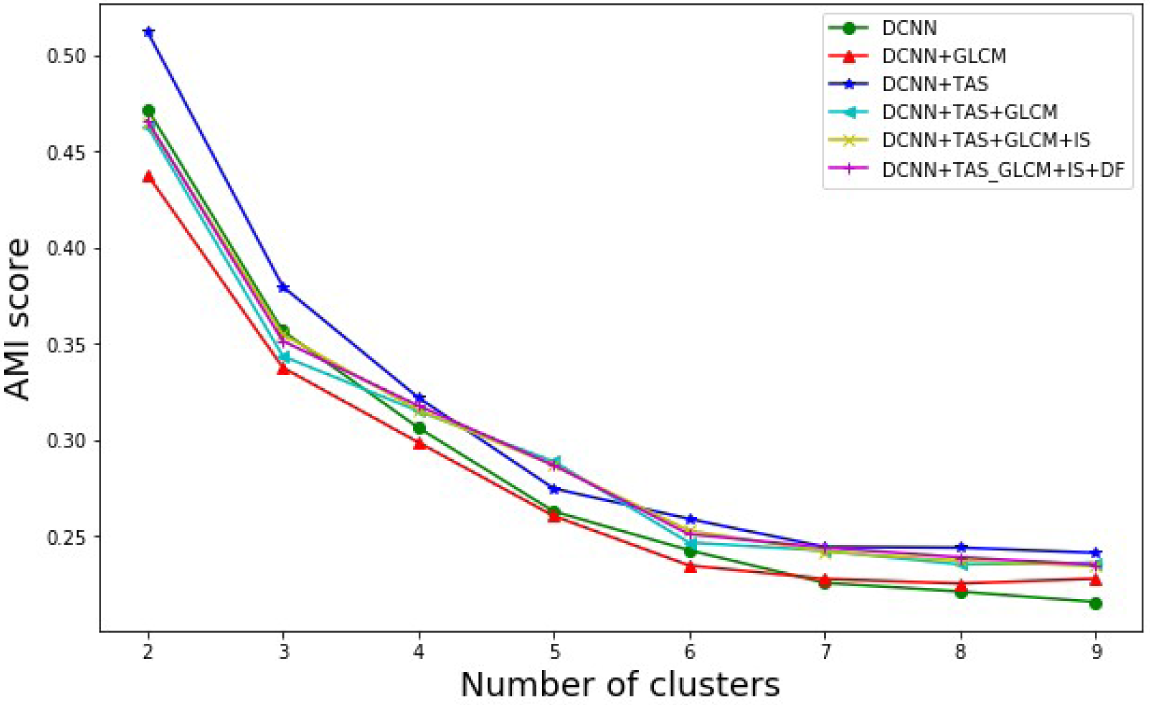
AMI score curves obtained with different combination of morphological features. 6 combinations have been investigated, which include DCNN only as the baseline, DCNN+GLCM, DCNN+TAS, DCNN+TAS+GLCM, DCNN+GLCM+IS, and DCNN+TAS+GLCM+IS+DF (see section 3.3 for the full names of the features). TAS features characterize the texture of an image and show the highest AMI score when concatenated with the DCNN features.

#### 4.1.3. Selecting clustering algorithm for MSI data

After optimizing the morphological feature set for H&E images, we used the segmentation results obtained using the morphological features as “pseudo ground truth” to evaluate different clustering algorithms for MSI data. AMI score was used again to measure the agreement between MSI-based clustering and the pseudo ground truth. KMeans is one the most commonly used clustering algorithms for MSI data. In practice, people often needs to determine which m/z variables are the key discriminants (molecular biomarkers) between clusters. However, KMeans algorithm doesn’t directly provide useful information to answer this question, so we have to resort to other techniques such as LDA. As mentioned in section 3.5, co-clustering algorithm associates each m/z variable to a pixel cluster id, which means it allows spatial subregions (pixel clusters) and m/z variables co-localized with each sub-region to be discovered simultaneously. Figure 4 (A) displays part of the original data matrix, where columns represent m/z variables, rows represent pixel ids, and value at each matrix element corresponds to the scaled abundance of certain m/z at certain pixel. The data matrix seems chaotic and it is hard to recognize any spectral pattern. However, after Co-Clustering, if we rearrange the data matrix by moving m/z variables and pixels of same cluster id to be contiguous, a chess board pattern appears as in Figure 4(B). It immediately becomes interpretable: m/z variables of the upper-left/lower-right square are more abundant for pixels of cluster 1/2 and, in a different perspective, pixels of the upper-left/lower-right square are clustered together because they have high abundance in a same group of m/z variables.

**Figure 4:**
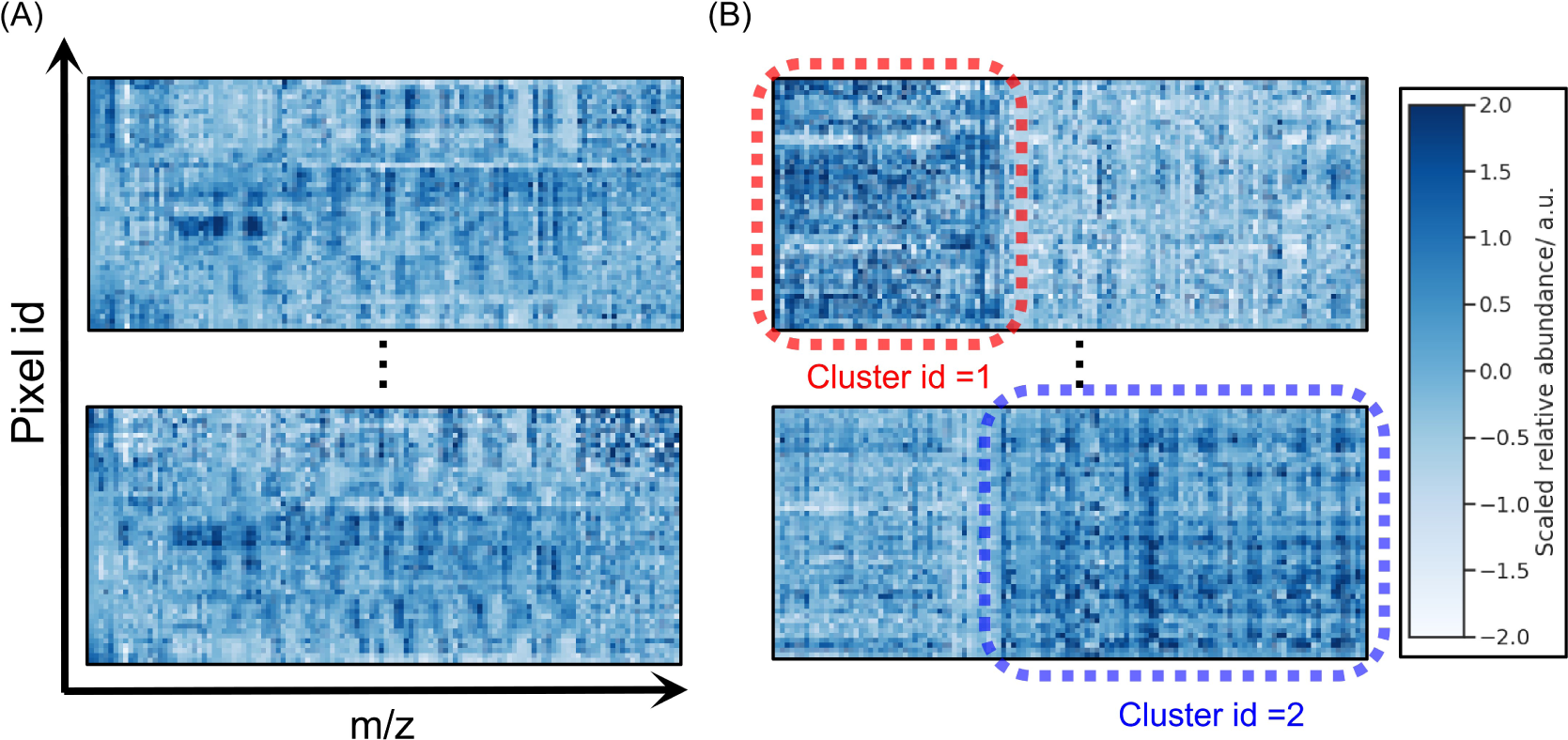
(A) Part of the original data matrix, columns corresponds to different m/z variables and rows correspond to different pixel ids. (B) new data matrix after co-clustering, where m/z variables and pixel ids are rearranged to move items of the same cluster contiguous.

MSI-based clusters obtained by KMeans and Co-Clustering respectively are compared in Figure 5 according to their AMI scores at different predefined cluster numbers K. Co-Clustering has a higher AMI score at K=2 and KMeans performs better at K=3 and K=4, while from K=5 to 9 they become nearly identical. The results indicate that Co-Clustering is competitive with KMeans and is a handier tool due to its simultaneous determination of potential biomakers. Figure 6(A) displays the mean mass spectrum of the kidney tissue sample, blue peaks are assigned to pixel cluster 1 and red peaks are assigned to pixel cluster 2. Figure 6(B) and (C) show the spatial distribution of “top” m/z variables associated to each of the clusters 1 and 2. [42] defines the top column variables as those whose internal edge weights in a bipartite graph are the greatest. Specifically in our case, the top m/z variable of a cluster is the one whose sum of scaled abundance over all pixels assigned to that cluster is the greatest. It should be observed that m/z 140.08 clearly co-localizes with pixel cluster 1 (which corresponds to pelvis),and m/z 738.52 co-localizes with the outer part of pixel cluster 2 (which corresponds to renal medulla).

**Figure 5:**
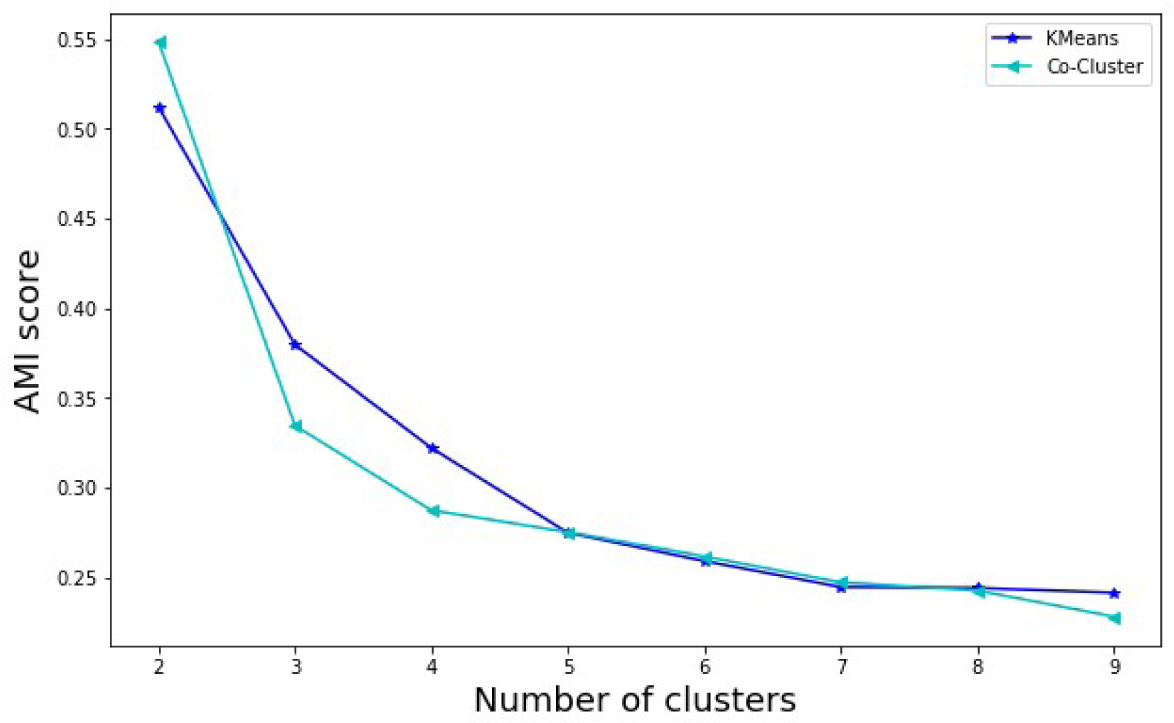
AMI score curves obtained with different clustering algorithms. Co-clustering algorithm shows a comparable performance with conventional KMeans algorithm.

**Figure 6:**
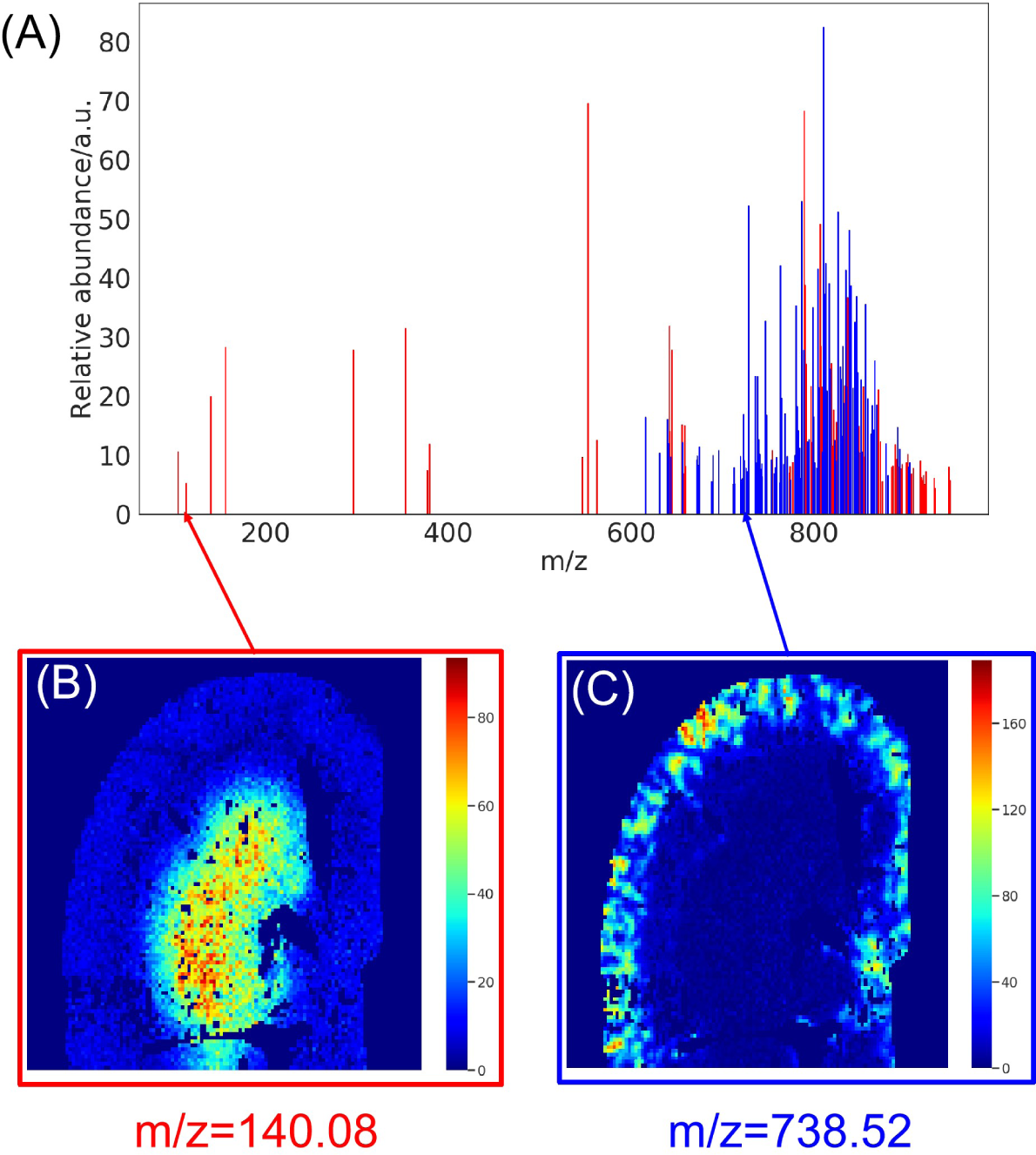
(A) The mean spectrum of all pixels with m/z peaks assigned to corresponding segments/clusters (red for cluster id 1 and blue for cluster id 2). (B) Abundance map of ion 140.08 m/z, which is the characteristic ion of segment 1. (C) Abundance map of ion 738.52 m/z, which is the characteristic ion of segment 2.

### 4.2. Integrated tissue segmentation

For a piece of tissue section, two correlated but not necessarily identical segmentation results can be obtained based on the MSI-derived m/z spectra data and the H&E image-derived morphological features data. In order to fully exploit the bio-information in the two independent data sources, their clustering analysis results should be integrated to produce a more informative and reliable segmentation of the tissue sample. The integration approach was proposed as follows: (1) each pixel on the 2D tissue section was assigned a false colour according to its MSI-based cluster id (this is a typical visualization scheme for the clustering results of MSI data); (2) for each pixel, if its MSI-based cluster id and H&E-based cluster id were not identical, the transparency of this pixel was set to 20%. The MSI-based clustering was done by Co-Clustering algorithm described in section 4.1.3 and the H&E-based clustering followed the DCNN+TAS protocol described in section 4.1.2. Before the integration process, the two sets of cluster ids had to be matched up so that the same ids corresponded to roughly the same regions on tissue section. By increasing the transparency of pixels that are assigned to different clusters based on different data source, we can visualize the confidence level of the clustering result: solid colour means two data sources have consensus, so we are more confident about the assignment, while transparent pixel means two data sources have dissensus, so we are less confident about the assignment. In Figure 7, integrated segmentation maps are shown for K of 2,3, and 4. In (A), pixels whose cluster id = 1 are assigned red, pixels whose cluster id = 2 are assigned blue. In (B), green is added to represent cluster id 3 and grey is added in (C) for cluster id 4. In agreement with Figure 7, the number of unconfident pixels dramatically increases as K rises from 2 to 4. When K=2, most of the pixels are confident ones, which means the tissue segmentation based on the spatial heterogeneity of bio-chemical composition (MSI) can be closely reproduced based on that of histologic morphology (H&E image). Such double authentication guarantees that it is the biologically meaningful structure that gets reflected by the segmentation rather than noise or artifacts during data collection process. A closer look at the unconfident pixels shows that they very often appear at the boundary between the red and blue regions, there are two possible explanations for this: (1) the transitional nature of boundary tissue, which makes it sort of a “mixture” of the two sub-regions and therefore harder to be segmented correctly; (2) the tissue sections used for MSI and H&E staining may not be strictly identical, so the mapping between MSI pixel and H&E patch is not perfectly accurate.

**Figure 7:**
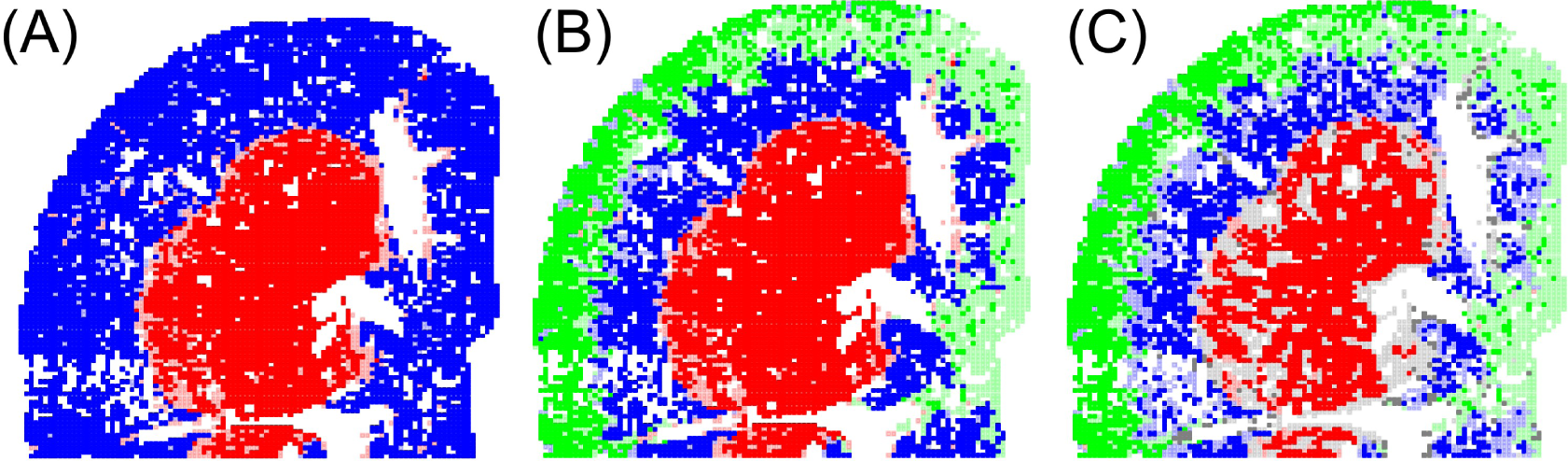
Integrated tissue segmentation map for (A) K = 2, (B) K = 3, and (C) K = 4. A pixel’s colour is solid if both MSI and morphological features assign the pixel to the same cluster id, and it becomes transparent if there is a disagreement. It is evident that when K = 2, the tissue segmentation is almost reproduced with the two independent bio-information sources, which strongly supports its validity.

## 5. Conclusion

We provided a more reliable approach to unsupervised tissue segmentation which exploited bio-information from two distinct data sources MSI and H&E image. By maximizing the match between MSI-based and H&E image-based clusterings, a morphological feature set, comprised of the pooling layer outputs of the fourth block of VGG16 and the TAS features, was established for histopathology image analysis. Similar approach was followed for the MSI data, which selected K=2 as the most probable number of clusters (i.e. number of sub-regions of the tissue sample) and compared different clustering algorithms. Co-Clustering algorithm was chosen due to its capability of automatically unearthing characteristic m/z variables associated with each sub-regions. For instance, ions with m/z = 140.08 colocalized with subregion pelvis and ions with m/z = 738.52 colocalized with subregion renal medulla. By integrating the segmentation outcomes on the basis of chemical information and morphological information, we evaluated the validity of the unsupervised segmentation and visualized the confidence level of cluster assignment for each location on the tissue section.

## References

[1] N. Verbeeck, R. M. Caprioli, R. V. de Plas, Unsupervised machine learning for exploratory data analysis in imaging mass spectrometry, Mass Spectrometry Reviews (2020).

[2] H. A. Vrooman, C. A. Cocosco, F. V. D. Lijn, R. Stokking, M. A. Ikram, M. W. Vernooij, M. M. B. Breteler, W. J. Niessen, Multi-spectral brain tissue segmentation using automatically trained k-nearest-neighbor classification, Neuroimage 37 (2007) 71–81.

[3] S. Wu, S. P. Weinstein, E. F. Conant, D. Kontos, Automated fibroglandular tissue segmentation and volumetric density estimation in breast mri using an atlas-aided fuzzy c-means method, Medical Physics 40 (2013).

[4] Liam, A., McDonnell, Ron, M.A., Heeren, Imaging mass spectrometry, Mass Spectrometry Reviews (2007).

[5] J. L. Norris, R. M. Caprioli, Imaging mass spectrometry: A new tool for pathology in a molecular age, Proteomics Clinical Applications 7 (2013) 733–738.

[6] K. Tanaka, H. Waki, Y. Ido, S. Akita, Y. Yoshida, T. Yoshida, T. Matsuo, Protein and polymer analyses up to m/z 100 000 by laser ionization time-of-flight mass spectrometry, Rapid communications in mass spectrometry 2 (1988) 151–153.

[7] Z. Takats, J. M. Wiseman, B. Gologan, R. G. Cooks, Mass spectrometry sampling under ambient conditions with desorption electrospray ionization, Science 306 (2004) 471–473.

[8] E. H. Seeley, R. M. Caprioli, Molecular imaging of proteins in tissues by mass spectrometry, Proceedings of the National Academy of Sciences of the United States of America 105 (2008) 18126–18131.

[9] P. Chaurand, S. A. Schwartz, R. M. Caprioli, Imaging mass spectrometry: a new tool to investigate the spatial organization of peptides and proteins in mammalian tissue sections., Current Opinion in Chemical Biology 6 (????) 676–681.

[10] D. Touboul, A. Brunelle, O. Laprévote, Mass spectrometry imaging: Towards a lipid microscope?, Biochimie 93 (2011) 0–119.

[11] Y. Sugiura, M. Setou, Imaging mass spectrometry for visualization of drug and endogenous metabolite distribution: Toward in situ pharmacometabolomes, J Neuroimmune Pharmacol 5 (2010) 31–43.

[12] P. Inglese, J. S. Mckenzie, A. Mroz, J. Kinross, K. Veselkov, E. Holmes, Z. Takats, J. K. Nicholson, R. C. Glen, Deep learning and 3d-desi imaging reveal the hidden metabolic heterogeneity of cancer, Chemical Science 8 (2017) 3500–3511.

[13] G. Mccombie, D. Staab, M. Stoeckli, R. Knochenmuss, Spatial and spectral correlations in maldi mass spectrometry images by clustering and multivariate analysis, Analytical Chemistry 77 (2005) 6118–6124.

[14] S. Deininger, M. Ebert, A. Futterer, M. Gerhard, C. Rocken, Maldi imaging combined with hierarchical clustering as a new tool for the interpretation of complex human cancers., Journal of Proteome Research 7 (2008) 5230–5236.

[15] T. Alexandrov, M. Becker, S. Deininger, G. Ernst, L. Wehder, M. Grasmair, F. Von Eggeling, H. Thiele, P. Maass, Spatial segmentation of imaging mass spectrometry data with edge-preserving image denoising and clustering., Journal of Proteome Research 9 (2010) 6535–6546.

[16] J. H. Kobarg, Efficient spatial segmentation of large imaging mass spectrometry datasets with spatially aware clustering, Bioinformatics 27 (2011) p.230–238.

[17] K. D. Bemis, A. Harry, L. S. Eberlin, C. R. Ferreira, O. Vitek, Probabilistic segmentation of mass spectrometry images helps select important ions and characterize confidence in the resulting segments, Molecular Cellular Proteomics Mcp 15 (2016) mcp.O115.053918.

[18] W. M. Abdelmoula, B. Balluff, S. Englert, J. Dijkstra, M. J. T. Reinders, A. Walch, L. A. Mcdonnell, B. P. F. Lelieveldt, Data-driven identification of prognostic tumor subpopulations using spatially mapped t-sne of mass spectrometry imaging data, Proceedings of the National Academy of Sciences of the United States of America 113 (2016) 12244–12249.

[19] M. N. Gurcan, L. E. Boucheron, A. Can, A. Madabhushi, N. M. Rajpoot, B. Yener, Histopathological image analysis: A review, IEEE Reviews in Biomedical Engineering 2 (2009) 147–171.

[20] F. A. Spanhol, L. S. Oliveira, C. Petitjean, L. Heutte, A dataset for breast cancer histopathological image classification, IEEE Transactions on Biomedical Engineering 63 (2016) 1455–1462.

[21] B. Kieffer, M. Babaie, S. Kalra, H. R. Tizhoosh, Convolutional neural networks for histopathology image classification: Training vs. using pretrained networks, in: 2017 Seventh International Conference on Image Processing Theory, Tools and Applications (IPTA), 2018.

[22] Z. Han, B. Wei, Y. Zheng, Y. Yin, K. Li, S. Li, Breast cancer multisclassification from histopathological images with structured deep learning model, Scientific Reports 7 (2017) 1–10.

[23] J. Barker, A. Hoogi, A. Depeursinge, D. L. Rubin, Automated classification of brain tumor type in whole-slide digital pathology images using local representative tiles, Medical Image Analysis 30 (2016) 60–71.

[24] T. P. Siegel, G. Hamm, J. Bunch, J. Cappell, J. S. Fletcher, K. Schwamborn, Mass spectrometry imaging and integration with other imaging modalities for greater molecular understanding of biological tissues, Molecular Imaging and Biology 20 (2018) 888–901.

[25] P. Mobadersany, S. Yousefi, M. Amgad, D. A. Gutman, J. S. BarnholtzSloan, J. E. V. Vega, D. J. Brat, L. A. Cooper, Predicting cancer outcomes from histology and genomics using convolutional networks, Proceedings of the National Academy of Sciences 115 (2018) E2970–E2979.

[26] R. J. Chen, M. Y. Lu, J. Wang, D. F. Williamson, S. J. Rodig, N. I. Lindeman, F. Mahmood, Pathomic fusion: An integrated framework for fusing histopathology and genomic features for cancer diagnosis and prognosis, arXiv preprint 1912.08937 (2019).

[27] A. Walch, S. Rauser, S.-O. Deininger, H. Höfler, Maldi imaging mass spectrometry for direct tissue analysis: a new frontier for molecular histology, Histochemistry and cell biology 130 (2008) 421.

[28] T. K. Sinha, S. Khatib-Shahidi, T. E. Yankeelov, K. Mapara, M. Ehtesham, D. S. Cornett, B. M. Dawant, R. M. Caprioli, J. C. Gore, Integrating spatially resolved three-dimensional maldi ims with in vivo magnetic resonance imaging, Nature Methods 5 (2007) 57–59.

[29] R. Van de Plas, J. Yang, J. Spraggins, R. M. Caprioli, Image fusion of mass spectrometry and microscopy: a multimodality paradigm for molecular tissue mapping, Nature methods 12 (2015) 366–372.

[30] F. Vollnhals, J.-N. Audinot, T. Wirtz, M. Mercier-Bonin, I. Fourquaux, B. Schroeppel, U. Kraushaar, V. Lev-Ram, M. H. Ellisman, S. Eswara, Correlative microscopy combining secondary ion mass spectrometry and electron microscopy: comparison of intensity–hue–saturation and laplacian pyramid methods for image fusion, Analytical chemistry 89 (2017) 10702–10710.

[31] E. Reinhard, M. Adhikhmin, B. Gooch, P. Shirley, Color transfer between images, IEEE Computer Graphics and Applications 21 (2001) 34–41.

[32] L. Cooper, Histomicstk, 2016. URL: https://github.com/DigitalSlideArchive/HistomicsTK.

[33] J. L. Norris, D. S. Cornett, J. A. Mobley, M. Andersson, E. H. Seeley, P. Chaurand, R. M. Caprioli, Processing maldi mass spectra to improve mass spectral direct tissue analysis, International Journal of Mass Spectrometry 260 (2007) 212–221.

[34] S.-O. Deininger, D. S. Cornett, R. Paape, M. Becker, C. Pineau, S. Rauser, A. Walch, E. Wolski, Normalization in maldi-tof imaging datasets of proteins: practical considerations, Analytical and Bioanalytical Chemistry (2011).

[35] Kyle, D., Bemis, April, Harry, Livia, S., Eberlin, Christina, F. and, Cardinal: an r package for statistical analysis of mass spectrometrybased imaging experiments: Fig. 1., Bioinformatics (2015).

[36] R. M. Haralick, Texture features for image classification, IEEE Trans Smc 3 (1973).

[37] N. A. Hamilton, R. S. Pantelic, K. Hanson, R. D. Teasdale, Fast automated cell phenotype image classification 8 (2007) 110–0.

[38] K. Weiss, T. M. Khoshgoftaar, D. D. Wang, A survey of transfer learning, Journal of Big Data 3 (2016) 9.

[39] Y. LeCun, Y. Bengio, G. Hinton, Deep learning, nature 521 (2015) 436–444.

[40] K. Simonyan, A. Zisserman, Very deep convolutional networks for large-scale image recognition, Computer Science (2014).

[41] M. D. Zeiler, R. Fergus, Visualizing and understanding convolutional networks, in: European conference on computer vision, Springer, 2014, pp. 818–833.

[42] I. S. Dhillon, Co-clustering documents and words using bipartite spectral graph partitioning, in: Proceedings of the seventh ACM SIGKDD international conference on Knowledge discovery and data mining, 2001, pp. 269–274.

[43] F. Pedregosa, G. Varoquaux, A. Gramfort, V. Michel, B. Thirion, O. Grisel, M. Blondel, P. Prettenhofer, R. Weiss, V. Dubourg, J. Vanderplas, A. Passos, D. Cournapeau, M. Brucher, M. Perrot, E. Duchesnay, Scikit-learn: Machine learning in Python, Journal of Machine Learning Research 12 (2011) 2825–2830.

[44] R. Mormont, P. Geurts, R. Maree, Comparison of deep transfer learning strategies for digital pathology, in: 2018 IEEE/CVF Conference on Computer Vision and Pattern Recognition Workshops (CVPRW), 2018.

